# Transitory Invasion Graphs

**DOI:** 10.64898/2026.11.24.768834

**Authors:** Germain Agazzi, Frank Pennekamp, Alexandre Mauroy, Frédérik De Laender

## Abstract

Invasion graphs map community assembly by linking communities that contain stable equilibria through successful invasions. By design, they leave aside transient communities that arise when species arrive before a community settles. Transitory invasion graphs (TIGs) complement this framework by recording every observed community, stable or transient, weighted by its frequency. In 500 simulated competitive communities of 5–25 species, assembly expanded through many transient communities, then contracted onto a few attractors. Competition limited the expansion: stronger interactions produced fewer communities but more transitions among them. The contraction reflected the attractor type: stable-equilibrium communities settled on the endpoints predicted by invasion graphs, whereas cyclic and chaotic communities kept turning over, most often through communities away from those endpoints. Repeated invasion attempts also rescued species with negative invasion growth rates and occasionally displaced uninvadable equilibria. Together, invasion graphs and TIGs reveal both where assembly can end and what is observed along the way.

## 1 Introduction

Community assembly describes how species colonizations and extinctions shape ecological communities through time. During assembly, species from a larger species pool arrive in a community through invasion attempts [1]. Each attempt introduces a tiny population of a potentially new species, after which this population either shrinks into oblivion or grows and affects established community members [2] and ecosystem functions [3, 4]. However, another species may arrive before either outcome occurs, leaving a transient community containing vanishing species. As species arrive and disappear, they alter resident species densities, community composition [2], and ecosystem functioning [3, 4].

Community assembly has been studied since the 19^th^ century [5] and is considered a key concept in ecology, as it explains why some species are present or absent in a given ecosystem, and how communities can change over time [6, 7]. Yet, the mechanisms underlying observed species successions remain elusive because tractable theory requires assumptions that do not necessarily match reality [8]. One core assumption is that assembly is viewed as a sequence of locally stable (though invadable) equilibria (in these paper, we will refer to these as “equilibrium/equilibria”); these are compositions that, without invasion, would persist indefinitely through time. A corollary is that invasions must be considered rare enough such that they only occur after a community has settled into such an equilibrium.

The invasion graph (IG), also called assembly graph, is one tool that leverages the assumption of stable communities to represent assembly as a network whose nodes are stable communities (communities containing an equilibrium), connected by successful invasions [6–10]. For example, in an IG, the node {1,2} represents the community with species 1 and 2. If there is an edge between the nodes {1,2} and {1,3}, this means that successful invasion by species 3 moves a community with species 1 and 2 to a community with species 1 and 3. This example illustrates how, because of the locally stable equilibrium assumption, the transient community {1,2,3} is absent from the IG, even though it would be observed empirically. This suggests that IGs overlook the many numerous transient communities [2, 11, 12], as well as invasions into them, and the diversity of associated assembly pathways [2].

Transient communities can also contribute to and shape long-term community and ecosystem dynamics even if they vanish [2, 11, 12]. For example, catalytic species are species that are present during the assembly process and essential to it, but go extinct after a long time, such as pioneer species [7, 8, 13]. Transient invaders can also have important consequences, such as shifts from one equilibrium to another [14]. Moreover, some of these transient communities can persist for a long time period [2, 11, 12], or even be locked into transient dynamical regimes by recurring invasions [2, 12], leading to chaotic dynamics [15]. The fact that the rate of invasion attempts can match the timescale of local dynamics [16, 17] further complicates the resulting dynamics, leading to more transient communities and assembly pathways.

Here, we propose a new quantitative approach to study community assembly that explicitly accounts for transient dynamics and high invasion rates. Specifically, we introduce the *transitory invasion graph (TIG)*, with nodes that represent communities that are stable (*sensu* above) *or* transient, and edges that represent a transition between two communities. It can be an observed invasion with invader density above a detection threshold (at any time), or an extinction when a species’ density falls below a detection threshold (Fig. 1 A). Moreover, the TIG integrates the whole assembly trajectory up to a selected time-point by weighting its nodes and edges according to their occurrence throughout history. This time-integration gives insight into the dominant communities at various moments throughout assembly history. We then apply our approach by simulating TIGs for random communities of 5 to 25 species that interact antagonistically (*e*.*g*. through competition), testing a wide range of interaction parameters. We showcase TIGs for canonical community dynamics: (1) a single and globally stable equilibrium, (2) multiple stable equilibria, depending on invasion history, (3) cycling, and (4) chaos.

**Figure 1:**
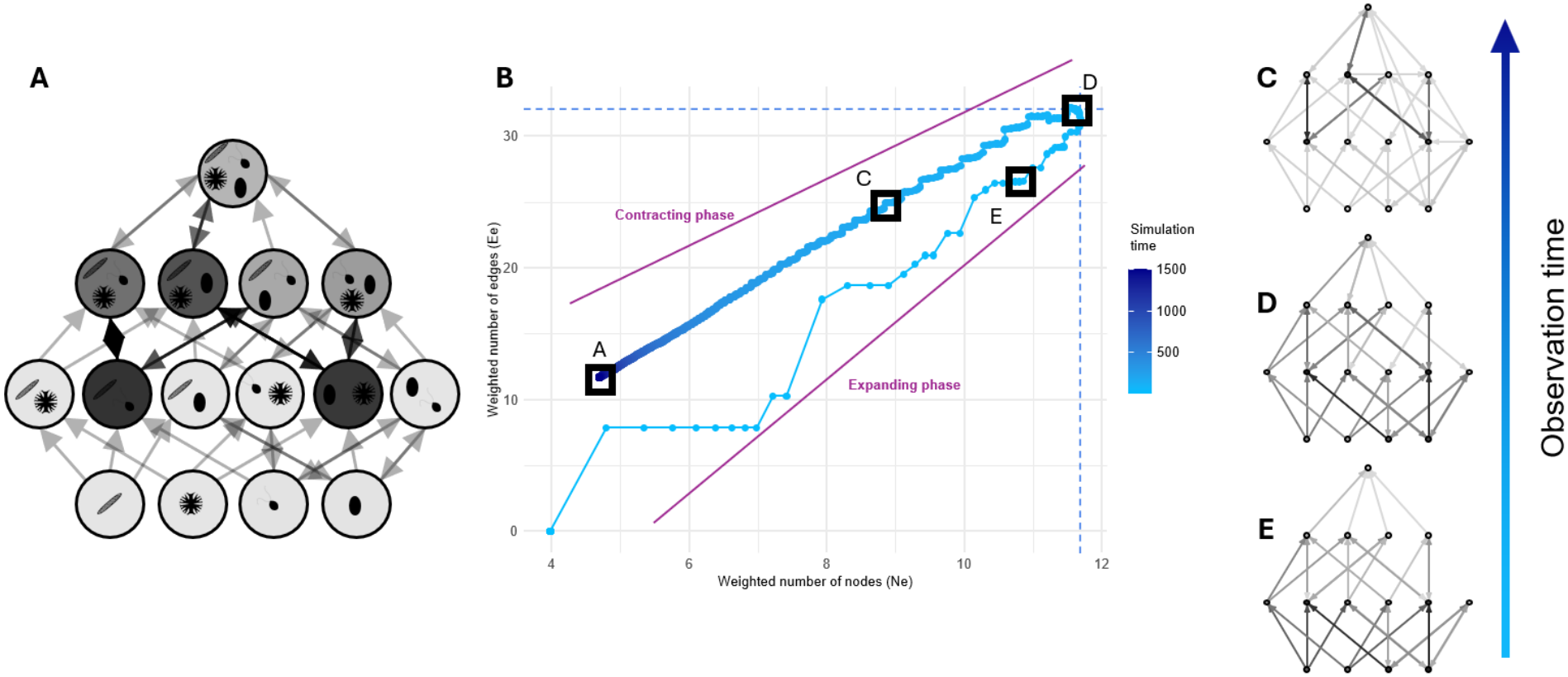
Example of a transitory invasion graph and its *E*_*e*_ *∼N*_*e*_ trajectory. (A) An example of a transitory invasion graph (TIG) with a species pool of *S* = 4 and assembly time *t* = 5000; darker nodes and edges have higher weights. (B) The *E*_*e*_ *∼N*_*e*_ trajectory of the same TIG; each dot corresponds to the TIG at a different time step. (C), (D), and (E) show the same TIG at different observation time, respectively after, at, and before the maximal *N*_*e*_ and *E*_*e*_ values. Squares in panel (B) correspond to the TIG in panels (A), (C), (D), and (E). Dashed blue lines correspond to maximal values of *N*_*e*_ and *E*_*e*_ and purple lines indicate the two phases.

The TIGs reveal that community assembly has a characteristic expansion–contraction structure, and that the shape of this trajectory contains information about the underlying ecological dynamics. More specifically: TIGs initially expand, as invasion order generates a diversity of transient communities and transitions, and subsequently contract, as dynamics increasingly concentrate around a smaller set of attractors. Interspecific interactions limit how far the TIG expands, whereas the nature of the attractors and continued turnover determine how it contracts and where it ends. These results show that TIGs help to quantify community assembly more completely than before.

## 2 Results

### 2.1 Transitory invasion graphs for random antagonistic communities

Transitory invasion graphs (TIGs) describe community assembly by tracking both the communities encountered during assembly and the transitions between them (Box 1). They are conceptually related to classical invasion graphs [6], but differ in important respects. First, their nodes are not restricted to communities containing a locally stable equilibrium, *i*.*e*. where each member species has a positive growth rate when rare in the community formed by the other members, but can include all observed communities, including transient ones. Second, their edges represent observed transitions between communities rather than the long-term consequences of invasion. Third, the structure of a TIG changes through time. To summarize the TIG structure throughout assembly history, we weigh nodes and edges by how often they occur through time. To this end, we use the effective numbers of nodes (*N*_*e*_) and edges (*E*_*e*_), calculated as the exponential of Shannon entropy (Eq. 4). These quantities measure the effective diversity of communities and transitions, respectively: they are high when many nodes or edges occur with similar frequencies, but approach the number of frequently occurring objects when a small subset dominates [18]. As illustrated by a motivating example, community assembly can be represented as a trajectory in *E*_*e*_ ∼ *N*_*e*_ space (Fig. 1 B).

Assembly typically starts with few observed communities and no observed transitions, and therefore low *N*_*e*_ and *E*_*e*_ = 0. This phase is characterized by many invasions of species that are bound to remain present for only a short time. The identity of those species depends on the invasion order. Random invasion orders therefore lead to random transient communities. This results in an accumulation of nodes, edges, and assembly pathways, leading to an increase in both *N*_*e*_ and *E*_*e*_ in an expanding phase of the TIG (Fig. 1 B, before square D). We summarize this accumulating phase by taking the maximum of *N*_*e*_ and *E*_*e*_ values across time. Whereas a TIG can reach its maximum *N*_*e*_ and *E*_*e*_ values at different times, these times are usually close: when the species pool size is small (*S* = 5), the median proportion of *N*_*e*_ relative to its maximum when *E*_*e*_ is at its maximum is 0.988, whereas the median proportion of *E*_*e*_ relative to its maximum when *N*_*e*_ is at its maximum is 0.981. In contrast, when the species pool size is larger, these proportions decrease (0.575 and 0.728 for *S* = 25), meaning that *N*_*e*_ reaches its maximum before *E*_*e*_ does (see supplementary information B).

#### Box1

**What is a transitory invasion graph (TIG)?**

TIGs are constructed from repeated observations of community assembly over a time interval *t*. Each node represents an observed community, defined by its combination of species, and nodes are organized into levels (*l*) according to species richness, ranging from 1 to *S*, where *S* is the size of the species pool. A directed edge links two communities and represents an observed transition between them . Edges can go from *l* to *l* + 1 upon invasion, from *l* to *l*− 1 upon extinction, and from *l* to *l* when both phenomena occur at the same time (species turnover).

Because the same community or transition can be observed repeatedly throughout the assembly process, nodes and edges are assigned weights corresponding to their frequencies of occurrence. At each time point, we therefore compute the effective numbers of nodes (*N*_*e*_) and edges (*E*_*e*_), see Methods section. Following *N*_*e*_ and *E*_*e*_ through time produces a trajectory in *E*_*e*_ ∼*N*_*e*_ space that describes how the diversity of communities and assembly pathways changes during assembly (Fig. 1 B).

As an informative benchmark, we consider a “null” TIG. This is a TIG originating from a species pool without interspecific interactions and with all intrinsic growth rates positive. This implies that every invasion by a new species is successful. Under no conditions is it possible to go from a node at level *l* to a node at level *l*− 1 on the null TIG. Instead, the probability of going from a node at level *l* to a node at level *l* + 1 is simply the probability that an invasion attempt involves a new species, *i*.*e*., 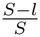. Considering this, it is possible to use Markov chains and edge probabilities to compute *P* (*t*) as a function of time *t* and frequency of invasion attempts *λ*, given the probability of being at each level *l* [19, 20]. Because all nodes and edges within a given level of a null TIG have the same weight, *P* (*t*) can be used to calculate the weight of a node and the weight of an edge at each level. The weight of a node associated with a level *l* is given by

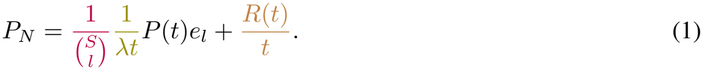

where *e*_*l*_ is the canonical unit vector selecting level *l*. The black expression gives the mean number of times level *l* is observed over the ⌊*λt*⌋ invasion attempts that occurred by time *t*. The green expression converts this number into the fraction of time spent at level, and the orange term is the time remaining after the last invasion attempt. The purple expression divides by the number of nodes at level to obtain the fraction of time spent at a single node. For an edge associated with a transition from level to level, we have

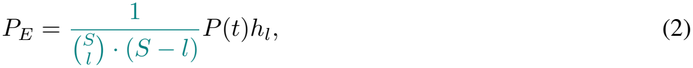

where 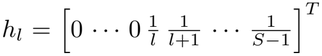 has its first *l* components equal to zero. The black expression gives the probability of observing a transition from level *l* to level *l* + 1. The blue expression divides this probability by the number of possible transitions from level *l*, giving the probability of observing one given transition. More details are given in suuplementary information A.

We simulated assembly across species pools of different size and different strengths of interspecific interaction, and analysed the expanding and contracting phases in all of these assembly. Across species pool sizes, the relationship between maximal node and edge values was well described by a power law (Fig. 2 A; *R*^2^ = 0.996). Interaction strength strongly modified this equation’s coefficients: strong interactions led to steeper relationships, suggesting that stronger antagonistic interactions reduce nodes and edges diversity. Higher interaction strength is associated with a lower maximum number of communities but more transitions between them. Furthermore, null TIGs, (*i*.*e*. TIGs for pools of species that don’t interact; Box 1), also reflected this power law, and provided a lower bound for the *E*_*e*_ ∼*N*_*e*_ relationship.

**Figure 2:**
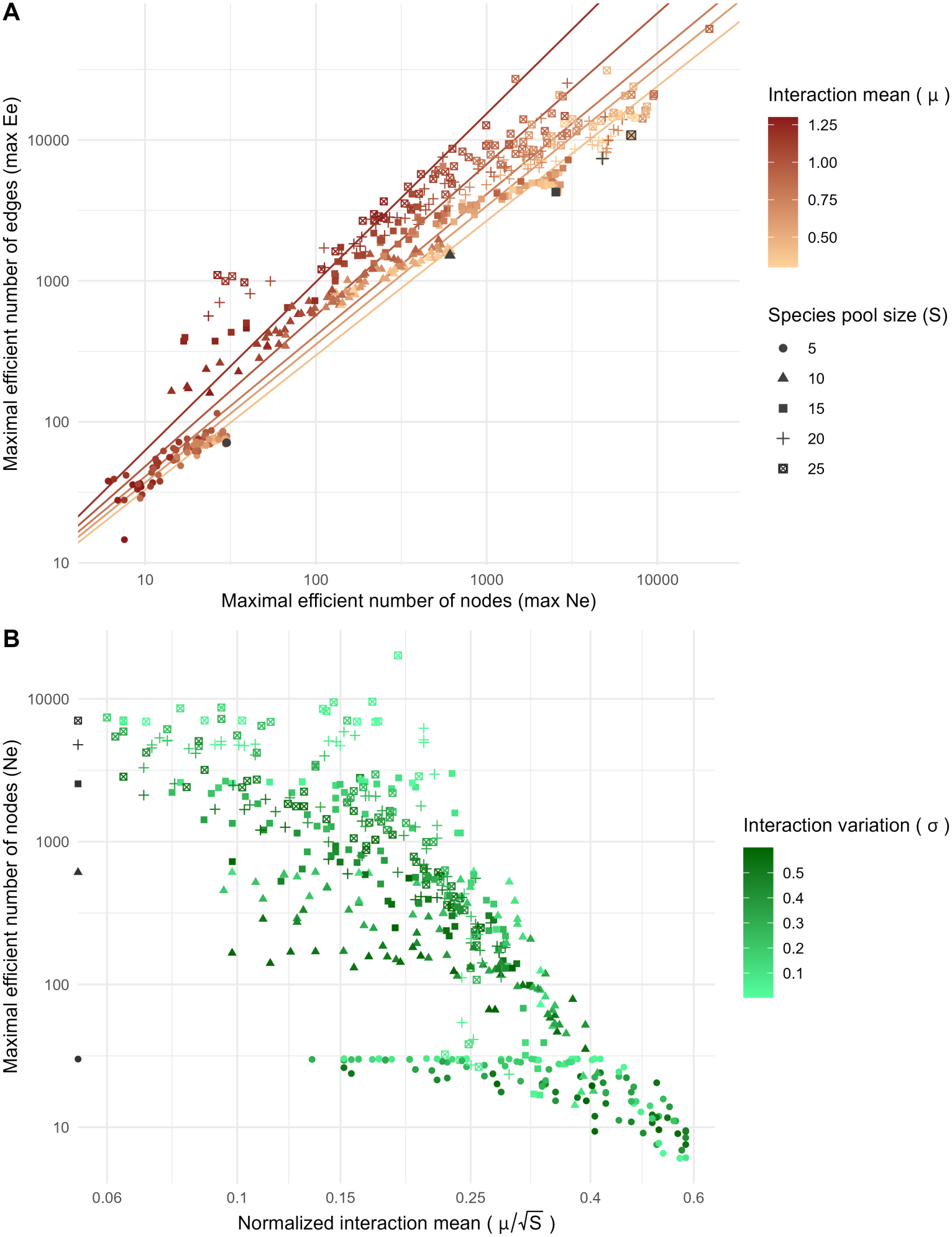
Position of the maximal *N*_*e*_ and *E*_*e*_ values of each TIG. (A) Position of the maximal *N*_*e*_ and *E*_*e*_ for each TIG in the log −log space. Colour indicates the interaction coefficient mean (*µ*), and shapes correspond to the species pool size (*S*). (B) Maximum *N*_*e*_ value as a function of 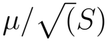. Colour indicates the interaction coefficient standard deviation (*σ*), and shape corresponds to *S*. Grey symbols correspond to null TIGs’ maximal values; their shapes correspond to *S*. For panel B, null TIGs’ dots are not placed at 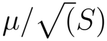 but just to the left of the plot for clarity.

Since the maxima of *E*_*e*_ and *N*_*e*_ are related, studying one of them suffices to understand how interspecific interactions determine the maximum extent of a TIG. Here, we consider *max*(*N*_*e*_). Because the interactions experienced by a species depend on the size of the species pool [21, 22], we scaled mean interspecific interaction strength *µ* by 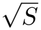.

We found that maximal *N*_*e*_ decreased as this scaled variable increased, meaning that TIGs become smaller, with fewer nodes, as interactions intensify. Also, the relationship between the maximal *N*_*e*_ and scaled mean interaction strength saturates in a way that depends on *S*. This reflects the fact that the number of communities in a TIG is capped by the number of available species (Fig. 2 B). This saturation is captured by the null TIG (Box 1), although the actual asymptote is typically lower than predicted by the null TIG, especially when interspecific interactions are strong, due to the pruning by species interactions.

After the expanding phase, once *max*(*N*_*e*_) is reached, competition increasingly drives the communities towards attractors (*i*.*e*., the states towards which densities converge in the long term), which reduces the number of communities and assembly pathways during the contracting phase. Because these attractors are determined by competition, they correspond to an uninvadable equilibrium. Indeed, we observed that densities were generally closer to such an equilibrium than randomly generated densities were. In our experiments, two exceptions were observed: (1) TIGs in which every reached node was an attractor, so that there was no contracting phase; and (2) TIGs without equilibria, in which the “least unstable” equilibrium (*i*.*e*., the one whose Jacobian eigenvalues had the smallest maximal real part, even when all of these values were positive) was the only attractor. We therefore always observed identifiable deterministic attractors from the perspective of local asymptotic stability.

From a trajectory point of view, this convergence leads to a reduction in *N*_*e*_ and *E*_*e*_, because successive time windows increasingly contain the same communities. The contracting phase therefore corresponds to an asymptotic decrease in *N*_*e*_ and *E*_*e*_ after they reach their maximal values. The contracting phase is thus identifiable as a general decrease along the trajectory.

During TIG contraction, *N*_*e*_ and *E*_*e*_ decrease at rates that may or may not be similar. If they are similar, the contracting phase of the trajectory is linear (both metrics are correlated); otherwise, it is curved. We measured this curvature by normalizing *N*_*e*_ and *E*_*e*_ during the contracting phase and then applying the Gini-like area index. This coefficient ranges from −0.5 (highly convex) to 0.5 (highly concave), with 0 corresponding to linear (see supplementary information C). We found that most trajectories were concave, indicating that *E*_*e*_ usually decreases more slowly than *N*_*e*_, meaning that the TIG reached attractor communities before the number of links (corresponding to transitions between communities) have settled. Moreover, curvature was correlated with transition probability (mean number of transition per time step) during the second half of assembly history (Fig. 3). A higher frequency results in more transitions, probably along the same edges, leading to greater edge weights and a faster decrease in *E*_*e*_, thus reducing the concavity of the contracting phase. TIGs with larger *S* had higher area index, showing that a perfect correlation between *N*_*e*_ and *E*_*e*_ is unlikely in species-rich communities.

**Figure 3:**
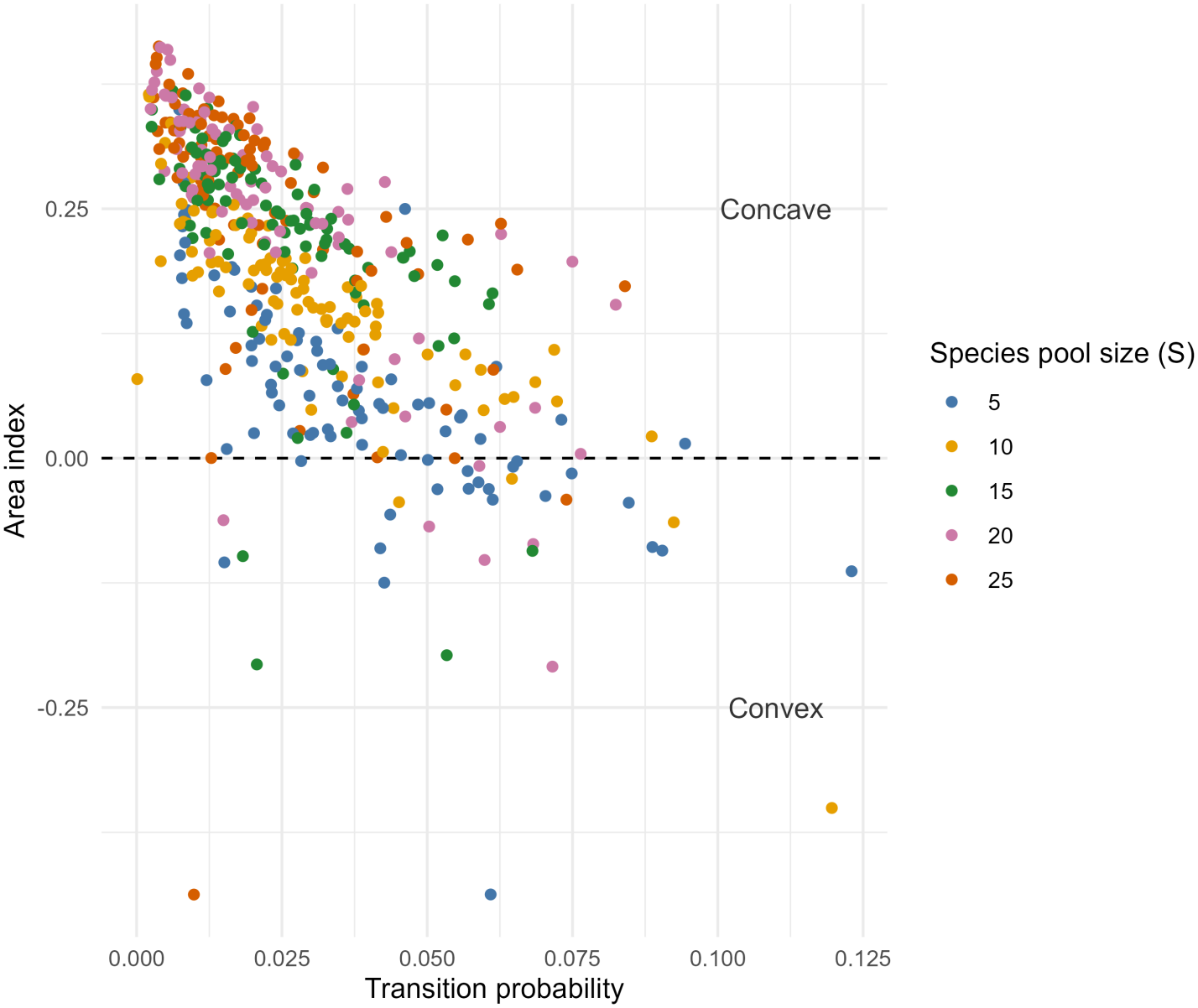
Relationship between the trajectory’s contracting phase curvature and transition probability. The y-axis corresponds to the area index, which is positive for concave contracting phases and negative for convex ones. The x-axis corresponds to the transition probability during the second half of the observation time. Colours indicate the species pool size *S*. Only TIGs with a transition probability *>* 0 during the second half of the assembly are shown in this plot.

In contrast to an invasion graph, the communities in a TIG at the end of assembly (when *N*_*e*_ and *E*_*e*_ have reached their final value) are not always communities containing an equilibrium, but can contain a persisting invader. This occurs when a species has a slightly negative invasion growth rate that is offset by repeated invasions (more details in the supplementary information D). This mainly occurs at high frequencies of invasion attempts (*λ*) and small species pool size (*S*). Indeed, if 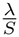 is high, each species attempts invasion more often, which increase its chance of persisting, despite a negative invasion growth rate.

Furthermore, the number of uninvadable equilibria that a species pool can produce is a lower bound for, rather than a predictor of, *N*_*e*_ at the end of assembly (Fig. 4). This can be explained by two phenomena: (1) *N*_*e*_ is not always stabilized at the end of our observations, and therefore does not necessarily represent the true final *N*_*e*_, and (2) some nodes can still be observed as transient communities even though they do not correspond to an equilibrium. However, some rare TIGs (around 1 out of 100) had an *N*_*e*_ lower than the number of equilibria. In these TIGs, a community containing an equilibrium at *l* = 1 (one species) could still be invaded due to repeated invasion attempts, leading to a transition to another equilibrium (see supplementary information D). This initial equilibrium is thus never maintained, leading to a lower number of nodes than expected from the number of uninvadable equilibria.

**Figure 4:**
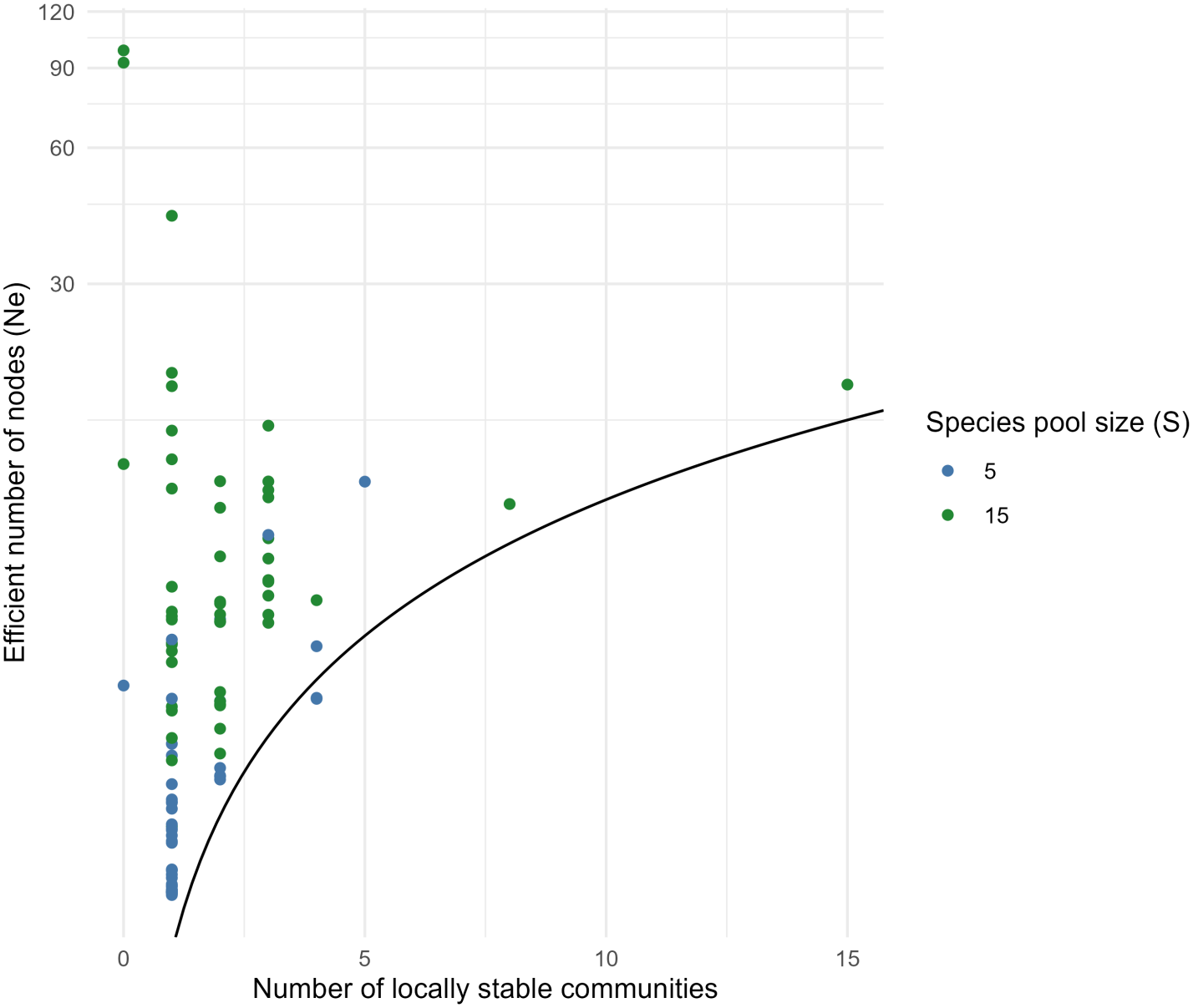
Effective number of nodes as a function of the number of uninvadable equilibria. Effective number of nodes (*N*_*e*_) at the end of the observations for 50 TIGs at *S* = 5 and 50 TIGs at *S* = 15 plotted against their number of communities containing an uninvadable equilibrium. The line represents the 1:1 relationship, and the y-axis is shown on a logarithmic scale.

### 2.2 Application to canonical community types

In this section, we apply TIGs to four contrasting community types, and explore how their different dynamics lead to different trajectories of the TIG’s two properties (*E*_*e*_, *N*_*e*_). These dynamics were (1) a single and globally stable equilibrium, (2) multiple equilibria, depending on invasion history, (3) cycling, and (4) chaos. Parameter settings are listed in the code (see supplementary information). For all of these TIGs, we recorded *N*_*e*_ and *E*_*e*_ at each time point to build their *E*_*e*_ ∼ *N*_*e*_ trajectories, which we then compared with those of a null TIG (Box 1).

We observed large differences among the trajectories of the four TIGs (Fig. 5). Only community type 1 matched the trajectory of the null TIG (based on a set of non-interacting species; Box 1), because invasions are mostly (type 1) or always (null TIG) successful and lead to a globally stable equilibrium in the community at *l* = *S*. For the other community types, TIGs followed trajectories that differed substantially from those of the null TIG.

**Figure 5:**
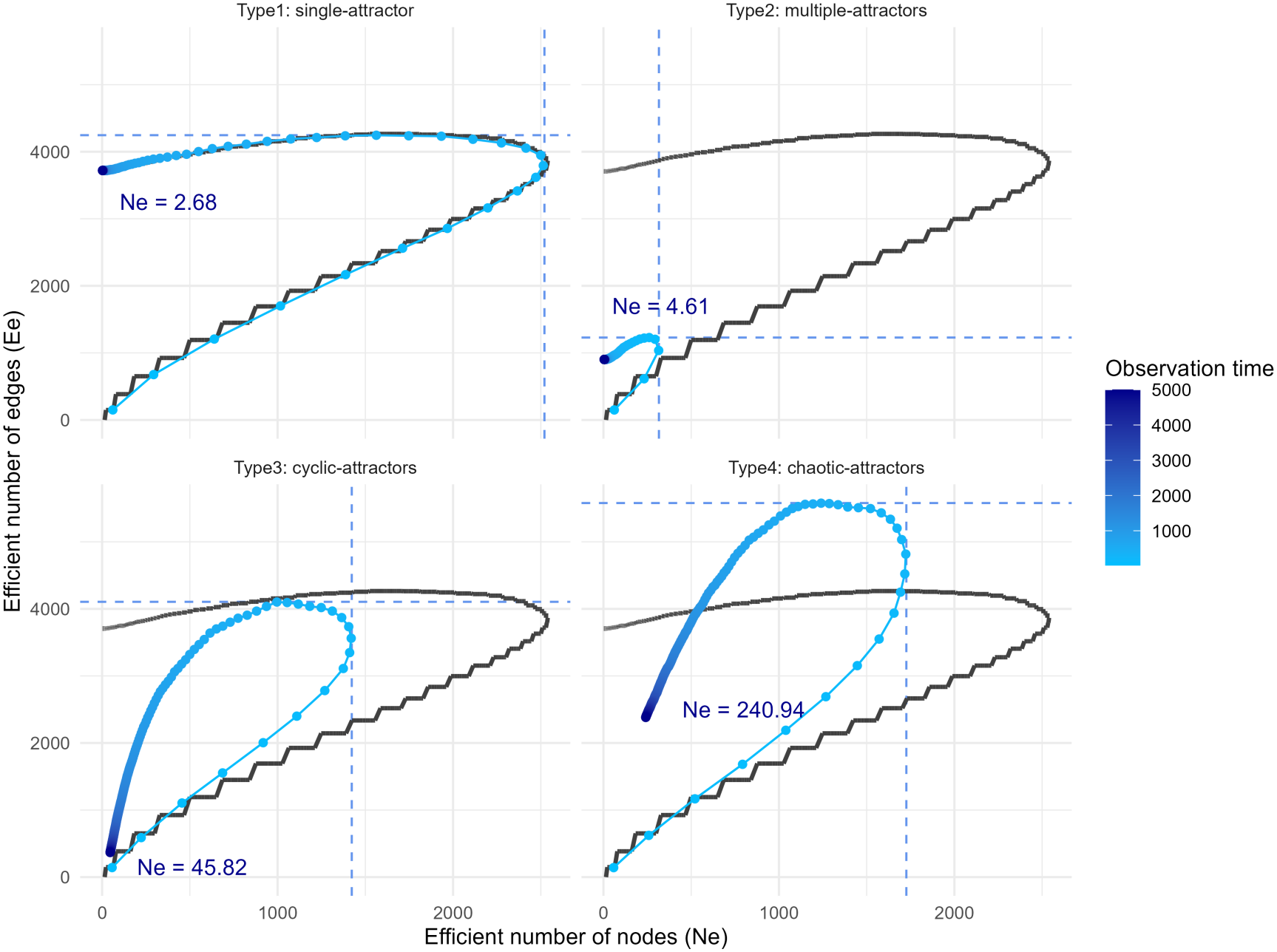
TIG trajectories for four canonical community types with *S* = 15. Each blue dot corresponds to the *N*_*e*_ and *E*_*e*_ values of a TIG at a given timepoint during assembly. The black line represents a null TIG (Box 1) with the same *S* and serves as a reference. Dashed blue lines correspond to maximal *N*_*e*_ and *E*_*e*_ values. Blue text corresponds to final *N*_*e*_ values.

The maximal values reached at the end of the expanding phase were lowest for community type 2. This is because interactions are stronger on average in this community type, causing transient communities with randomly arrived species to break up faster. Indeed, this TIG did not go beyond the seventh level (*l* = 7, communities with seven species) out of the 15 levels (its attractors were located at *l* = 5). Community types 3 and 4 were intermediate cases, reaching *l* = 11 and *l* = 13, whereas type 1 reached *l* = 15, as its attractor is located at this level (see supplementary information E).

In the contracting phase, the TIGs of community types 1 and 2 reduced *E*_*e*_ less than the other TIGs did, relative to their maximum values. This is because these two TIGs had very strong attractors, preventing successful invasions once these attractors were reached, which meant that edges gained very little weight (or none at all) over time. In contrast, community types 3 and 4, with cyclic and chaotic attractors, continued to exhibit changes in species densities and community composition, so some edges kept gaining weight, thereby affecting *E*_*e*_.

The final *N*_*e*_ for community types that exhibited fluctuating dynamics (3 and 4) was higher than that of those that did not (1 and 2). This was especially pronounced in the case of chaotic dynamics (community type 4), but also occurred in type 3. This is due to the large number of possible communities in the chaotic TIG, which increases the number of nodes and thus increases *N*_*e*_. Moreover, final *N*_*e*_ values were reached more quickly for communities that did not exhibit fluctuating dynamics (1 and 2) than for those that did (3 and 4).

In addition, the single-attractor and multiple-attractors TIGs had a transition probability of 0 during the second half of their observation time. This is visible in the flat aspect of the contracting phase, as explained previously. Moreover, during the assembly, the cyclic-attractors TIG had a lower transition probability (0.02) than the chaotic-attractors TIG (0.06), meaning that more transitions were observed over a same amount of time. That is also reflected in the more convex curvature of the cyclic attractors than for the chaotic attractors. The shape and values of the trajectories are thus good indicators of complexity and transition probability in these four typical TIGs.

We then asked what additional information a TIG provides compared with the classical invasion graph (IG) of the same community. The two graphs encode different information. An edge in an IG represents the outcome of an invasion into an equilibrium, and is independent of the observation time (*t*) and of the frequency of invasion attempts relative to the species pool size 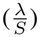. An edge in a TIG instead represents the probability of observing a transition, and therefore depends on both *t* and 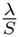. As a result, a TIG adds information on how often communities and transitions are observed during assembly, including after the communities has reached its attractors. For community types 1 and 2, this additional information was consistent with the IG: both graphs led to communities with no further transitions (uninvadable equilibria), and the TIG further indicated how frequently each of these communities was observed. For community types 3 and 4, the TIG revealed transitions that continued throughout the assembly, whereas the corresponding IGs converged to uninvadable communities. In these cases, the TIG provided a different description of long-term assembly.

Furthermore, TIGs captured only a fraction of IG nodes. Indeed, TIGs reached 11.30%, 2.47%, 9.87%, and 16.40% of IG nodes for TIG types 1 to 4, respectively, whereas all TIG nodes were also present in the IGs. This shows that observated assembly does not capture all the possible communities according to an IG, suggesting that IGs can reveal communities that may not be reachable when species invade transient communities. However, this metric is strongly affected by *t*. For example, null TIGs are expected to visit every node, as does the corresponding IG, but if *t* is too short, the TIG may miss some nodes, whereas the IG is not affected by *t* and always reaches every node.

Moreover, for some canonical communities, long-term assembly can differ between TIGs and IGs. Each IG converged to one or several nodes with incoming edges and no outgoing edges. These nodes correspond to communities that can be reached but not invaded (uninvadable equilibrium). Thus, if invasions do not occur into transient communities, the ecosystem converges to and remains in these communities over long times, with no further transitions. We observed this same pattern for types 1 and 2 (single or multiple attractors), but not for types 3 and 4 (cyclic or chaotic attractors). The first two TIGs showed that, regardless of the timing of invasion attempts, the ecosystem can converge to communities that remain uninvadable. This means that no further transitions occur, so edge weights become fixed, resulting in a flat contracting phase (Fig. 5, top). Moreover, both graphs converge to the same nodes in the IG and the TIG, with higher weights for these nodes in the TIG (Fig. 6, top).

**Figure 6:**
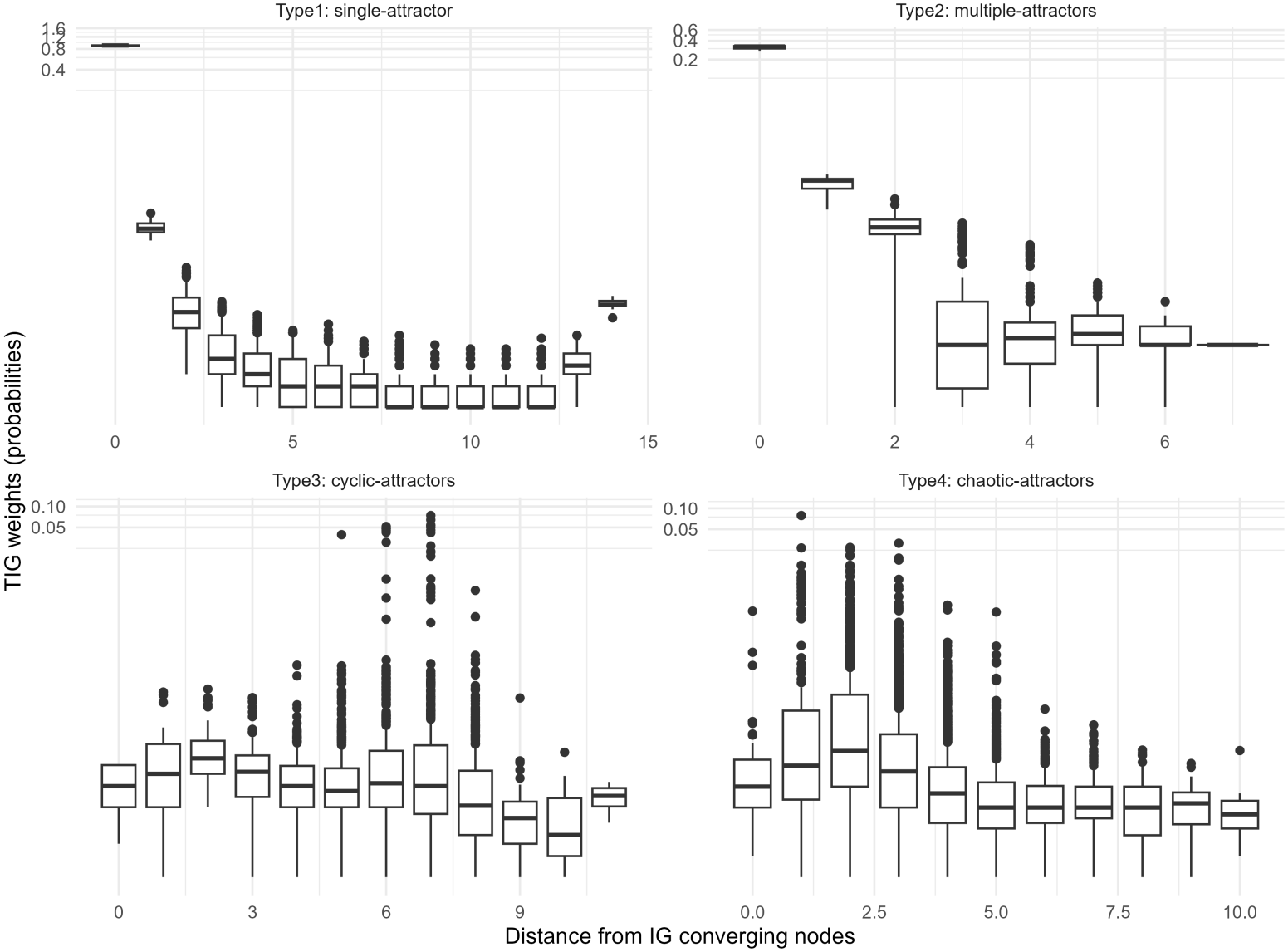
TIG nodes’ weight (as the probability of observing such node during the obser) as a function of their distance to the nearest IG converging node. Nodes at distance of 0 are the converging nodes themselves.

For types 3 and 4, IGs and TIGs lead to different interpretations. IGs suggest that, as for types 1 and 2, assembly converges to uninvadable communities. In TIGs, however, assembly converges to certain communities, but these keep changing with cyclic or chaotic dynamics: transitions continue to occur and gain weight, which reduces *E*_*e*_ values in the contracting phase (Fig. 5, bottom). Moreover, for types 3 and 4, TIG node weights do not correspond to the converging IG nodes (Fig. 6, bottom), showing that the most frequent communities are not those expected from the IG. This result confirms that the TIG protocol offers a different and flexible perspective on assembly processes, which may or may not match the one provided by IGs.

## 3 Materials and methods

### 3.1 Simulation assembly

To simulate assembly, we used the generalized Lotka-Volterra model described by equation 3:

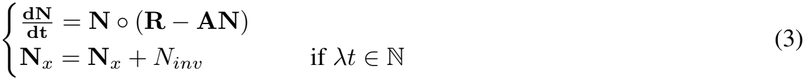

where **N** are species densities, **R** are intrinsic growth rates, **A** is the interaction matrix (higher values indicate stronger antagonistic interactions) [8, 23]. **N**_*x*_ is the density of species *x* (drawn at random from the *S* species), *λ* is the frequency of invasion attempts (for example, *λ* = 0.1 means one invasion attempt every 10 time steps), and *N*_*inv*_ is the invasion density. More specifically, we first fixed a pool of *S* species, and then started the simulation with a species randomly drawn from the pool at low density (*N*_*inv*_). Next, we added randomly chosen species (at the same low density *N*_*inv*_) at regular intervals of 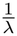 time steps. The invasion sequence is random: a given species may make several invasion attempts, or one, or none.

We considered a species as present (*i*.*e*. its invasion successful) when its density exceeded the invasion density (whatever the time), thereby defining a detection threshold. Furthermore, to reduce computation time, we put an extinction threshold where species with a density *N <* 0.0001 were set to *N* = 0. This procedure corresponds to a single assembly run. To build a transitory invasion graph (TIG), we ran an ensemble of simulations representing multiple invasion attempts and arrival sequences.

### 3.2 Simulating a TIG

We simulated 500 TIGs with species pool sizes (*S*) randomly selected from 5, 10, 15, 20, and 25. We ran longer simulations when *S* was large (to capture most of the assembly patterns, exact durations are given in the supplementary information F). For each species, we set all **R** values and all intraspecific interactions (the diagonal of **A**) to a standard value of 1 (thereby fixing the carrying capacity in the absence of interspecific interactions at 1). Interspecific interactions were sampled from normal distributions with means (*µ*) between 0.3 and 1.3, and standard deviations (*σ*) between 0.001 and 0.6. We set the invasion density to *N*_*inv*_ = 0.01 (1% of the carrying capacity in the absence of interspecific interactions) and thus the detection threshold to *N* = 0.0101. The extinction threshold was set to *N* = 0.0001.

During simulations, we recorded which species were present or absent to characterize the community. We then recorded the transitions between communities. In the graph, each community was represented as a node and each transition as a directed edge. Each node and edge was weighted by the probability of observing it during the simulations (defined as the number of occurrences of a given node or edge divided by the total number of node or edge occurrences). A more frequent node or edge thus had a higher weight (Fig. 1 A). We used these weights to compute the effective number of nodes (*N*_*e*_) and edges (*E*_*e*_) as the exponential of the Shannon index (*i*.*e*., the Hill number of order one [24]):

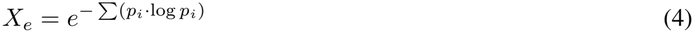

where *X*_*w*_ is either *N*_*e*_ or *E*_*e*_, and *p*_*i*_ is the probability of node or edge *i*. This metric returns a value close to the number of highly weighted objects when weight differences are large, and close to the total number of objects when weight differences are small.

### 3.3 Characterizing assembly with a TIG

For each TIG, we measured *N*_*e*_ and *E*_*e*_ at each time step and plotted them together as *E*_*e*_ ∼*N*_*e*_ trajectories (Fig. 1 B). These trajectories represent the node and edge diversity over time across all simulations of a single TIG. To characterize them, we extracted for each TIG the maximum *N*_*e*_ and *E*_*e*_ values, which represent the upper limits of the *E*_*e*_ ∼ *N*_*e*_ trajectories. Using these maximum values, we tested the relationship *max*(*E*_*e*_) = *a* · *max*(*N*_*e*_)^*b*^ by fitting the linear model log (*max*(*E*_*e*_)) ∼ log (*max*(*N*_*e*_)), where the slope is *b* and the intercept is log(*a*).

We also analyzed the curvature of the contracting phase (the end of the simulations) of these trajectories (see Results). To do so, we normalized the *N*_*e*_ and *E*_*e*_ values of this phase for each TIG to place them on the same scale (between 0 and 1). We then computed a Gini-like area index, defined here as the normalized area under the curve minus 0.5 [18]. This index is positive (maximum 0.5) when the curve is concave (∩ shape), negative (minimum −0.5) when it is convex (∪ shape), and zero when it is linear. The area under the curve was estimated by Monte Carlo integration [25]. The normalized contracting phases and their area index are provided in the supplementary information C. We compared this measure with the transition probability of the TIGs during the second half of the simulations. To do so, we computed, for each TIG, the probability of observing a transition at a randomly chosen time step.

To refine our analysis, we supplemented these 500 TIGs with 100 additional TIGs for stability analyses: 50 at *S* = 5 and 50 at *S* = 15. All other parameters and simulation settings were identical to those described above, except that we recorded densities (without detection thresholds) rather than communities. From these simulations, we also saved the interaction matrices in order to compute equilibria and assess their local asymptotic stability from the eigenvalues of the Jacobian matrix [23]. We then compared the observed densities with the densities of locally asymptotically stable equilibria (referenced in this paper as “equilibrium/equilibria”). Moreover, for each of these TIGs, we also generated random densities between 0 and 1 at each time step (the carrying capacity in the absence of interspecific interaction) to provide a baseline for the stability analyses. We did not compute *N*_*e*_ and *E*_*e*_ values for these 100 TIGs.

### 3.4 Application to canonical communities

Finally, we applied the TIG framework to communities with fixed interaction matrices. We first ran simulations to build “null TIGs”, in which all interspecific interactions were set to 0. We built one TIG for each value of *S* among 5, 10, 15, 20, and 25. The rest of the protocol (invasion attempt frequency, thresholds, etc.) remained unchanged. We also computed *N*_*e*_ and *E*_*e*_ to build *E*_*e*_ ∼ *N*_*e*_ trajectories. Furthermore, we calculated the predicted *N*_*e*_ and *E*_*e*_ values for null TIGs with a number of simulations tending to ∞ by using Markov chains [19, 20].

Lastly, we built TIGs for four canonical communities at *S* = 15: single-attractor, multiple-attractor, cyclic-attractor, and chaotic-attractor community. We applied the same protocol as described above and computed *N*_*e*_ and *E*_*e*_ to build *E*_*e*_∼ *N*_*e*_ trajectories. We also built classical invasion graphs (IG) for these four canonical communities. To do so, we identified all communities with a locally stable equilibrium (according to Jacobian matrix), whether invadable or not, and the outcome of every possible invasion into them. We then represented the results as a network in which nodes represent locally stable communities and edges represent invasion outcomes [7]. We compared the conclusions drawn from TIGs and IGs by observing reached nodes, dynamics, and transitions. We also defined “converging nodes” in IGs as nodes with incoming edges (reachable), but no outgoing edges (uninvadable). Lastly, we observed the relationship between the weight of TIG nodes and their distance to the nearest converging nodes.

## 4 Discussion

A central aim of community ecology is to understand how species interactions and the history of species arrivals together determine community composition. Current approaches, notably invasion graphs (IGs), describe assembly as a sequence of communities that each contain a stable equilibrium, connected by successful invasions. This description requires invasions to be rare enough for communities to settle between arrivals. As a result, it does not account for transient communities, although these are frequently observed [2, 11, 12] and can influence assembly even when invasions are rare [7]. Here, we introduced transitory invasion graphs (TIGs), an approach that describes assembly from all observed communities, whether stable or transient, and from the transitions between them, each weighted by how often it occurs. Because TIGs are built from observed assembly rather than from equilibrium analysis, they can be applied when invasion attempts are frequent, when dynamics fluctuate, and when species pools are too large for every subcommunity to be analyzed. Applied to simulated competitive communities, TIGs showed that assembly generally expanded through many transient communities before contracting onto a smaller set. The extent of the expansion depended on interaction strength, and the shape of the contraction depended on the type of attractor.

By observing the diversity of nodes and edges, we find that community assembly has a characteristic expansion–contraction structure in all simulated communities. This pattern suggests that, at the onset of assembly, a wide diversity of community compositions can be observed, but that this diversity decreases over time towards a handful of communities.

We interpret this observation as a signature of assembly stochasticity, which is therefore stronger in the early stages of assembly, whereas convergence towards the communities observed later is more deterministic. A similar pattern has been observed in the assembly of micro-organisms [26, 27] and macro-organisms [13], with an increased number of transient communities at the beginning of an assembly, *i*.*e*. during the expanding phase.

While the dynamic nature of the TIG implies that its structure (its nodes and links) is time-dependent, it still has a maximal effective number of nodes and edges. We found that these two maximal values were related across species pool sizes by a power law, a classic relationship in network science (Fig. 2 A). Ecologically, this implies that, for a fixed interspecific interaction strength, changing the species pool size does not strongly change how the TIG’s maximal node diversity scales with its maximal link diversity [28, 29]. Therefore, the processes driving the maximal node-link diversity relationship seem to be mainly affected by interaction strength rather than by species pool size.

After the expanding phase, deterministic population dynamics increasingly constrain assembly to a smaller set of communities associated with the TIG’s long-term attractors. Importantly, however, this does not imply that assembly reaches a fixed endpoint. Invasions continue to occur, so that even after long assembly times, stochastic invasion events can move the ecosystem through transient communities, especially in communities with fluctuating dynamics (*e*.*g*. Fig. 5, bottom) where observed communities change over time [12, 30]. The identity of the community observed at any given time can therefore remain partly contingent on recent invasion history, even when the underlying deterministic dynamics strongly constrain the set of reachable communities. Furthermore, it is already known that under the same conditions, communities can differ in the long term depending on invasion history, even though observed communities are usually influenced by the nearest equilibrium [2]. Assembly thus shifts from a broad stochastic exploration during the expanding phase to stochastic movement within a much more restricted region during the contracting phase, set by the number of stable communities acting as attractors [6,8]. Consequently, the effective number of TIG nodes decreases and eventually stabilizes around a restricted effective number of communities, rather than necessarily converging to a single, unchanging endpoint.

Nevertheless, the contracting phase can differ between TIGs. Some of them remain on attractor communities without further transitions, while others converge towards attractor communities with either few or many transitions. The first case corresponds to TIGs with strong attractors that prevent any further transitions [23, 31]. In this case, the effective number of edges will remain high (Fig. 5, top), making it possible to distinguish communities with strong attractors from communities without them. For communities without a strong attractor, transitions continue to occur [17, 27, 31]. These may be transitions from new communities directly to attractor communities; in this case, node and edge diversity are correlated. However, there are also transitions between previously observed communities that slowly converge towards attractor communities. In this second case, node and edge diversity are not correlated. Furthermore, long-term transitions probabilities depend on the species involved [31]: some communities will not undergo new invasions, while others may, leading to states where several communities are possible, and transitions can occur from one community to another, as observed with Markov chain models [32].

Moreover, we observed that IGs, compared to TIGs, will not capture some assembly patterns when the frequency of invasion attempts is high. On the other hand, TIGs did not represent as many communities as IGs, but focused on the most frequently observed ones, which are not necessarily the same [33]. Conversely, if the frequency of invasion attempts is low, communities have time to reach an equilibrium before the next invasion [6–8]; the TIG structure should thus be closer to the corresponding IG. Nevertheless, TIGs also depend on the observation time: node and edge weights represent the probability of observing the corresponding event during the observed time, whereas IG edges are time-independent [6–9]. Moreover, TIGs highlight frequently observed communities and transitions, whereas IGs do not [6, 7]. This illustrates that a TIG with a low invasion frequency is not equivalent to an IG. Because TIGs, unlike IGs, do not require communities to reach an equilibrium between invasion attempts, they make it possible to study the effects of key factors such as a high frequency of invasion attempts [7, 8, 34]. For example, if the frequency is low, the arrival order of species will matter more than when it is high [8, 16] (*i*.*e*., priority effects [2, 9]). IGs capture priority effects only in the first case, whereas TIGs can also be used to quantify how they change with the frequency of invasion attempts. Also, a high number of invasion attempts by the same species into an equilibrium can lead to a transition towards another community. This pattern can be compared to alien species invasions, which are strongly affected by “propagule pressure”, *i*.*e*. the density and frequency of invasion attempts. Furthermore, it has been observed that, for a low density of invasion attempts (as in our experiments), the effect of invasion attempt frequency on propagule was stronger, so that repeated invasion attempts by a species can lead to a higher probability of invasion success [35].

Together, these results suggest that key community properties could be inferred from its TIG trajectory. The extent of the expansion phase reflected the strength of interspecific interactions, the shape of the contraction phase reflected the type of attractor and the frequency of transitions, and the final effective number of nodes was generally bounded from below by the number of uninvadable equilibria. However, the generality of these relationships remains to be established, as our experiments were conducted under a restricted set of conditions. First, we held the frequency of invasion attempts constant; systematically varying this frequency would help determine its influence on TIG trajectories. Second, we considered only competitive interactions within the generalized Lotka-Volterra framework, excluding other interaction types such as predation, which can exhibit different cyclic dynamics [30] and potentially alter TIG structure. Third, we assumed constant environmental conditions throughout the assembly, whereas natural environments are typically variable. Whether environmental change would cause a community to shift between distinct TIGs, or instead give rise to a new TIG structure shaped by the history of environmental conditions, remains an open question.

We introduced transitory invasion graphs (TIGs), a framework for describing community assembly that accounts for all observed communities, whether stable or transient, rather than focusing exclusively on equilibria. Applied to simulated competitive communities, TIGs revealed two general phases of assembly: an expansion phase, during which the random order of species arrivals generates a large number of transient communities, followed by a contraction phase, during which dynamics converge towards a smaller set of attractor communities without necessarily reaching a fixed endpoint. In the canonical communities we examined, TIGs converged to the same endpoints as invasion graphs (IGs) when dynamics settled at stable equilibria. However, under cyclic or chaotic dynamics, the most frequently observed communities differed from these endpoints. Moreover, because TIGs include only communities that are actually reached during assembly, they avoid the need to analyze every possible subcommunity of the species pool and should therefore be applicable to larger species pools than IGs.

Because TIGs rely on observed community compositions rather than equilibrium analysis, they could also be constructed from replicated time series of community composition, such as those obtained from microcosms [7] or microbiomes [36]. This would enable comparisons of assembly across experimental and natural communities, particularly those characterized by frequent invasions and widespread transient communities [12]. TIGs could thus also prove useful in contexts where community assembly is actively managed, including ecosystem management [9, 12] and microbiome management [9, 36].

