## supplementary material for "Transitory Invasion Graphs"

### Supplementary information

This appendix summarises supplementary information from the **Transitory Invasion Graphs** article. In this article, we developed a new tool to study community assembly: the transitory invasion graph (TIG). Every data were generated through modelling, and datasets and R code can be obtained by contacting **.

#### A Null TIG $P_N$ and $P_E$

We first imagined a "null TIG" with no interspecific interactions, in which we can mathematically demonstrate the weights for nodes and edges. These were used to measure at every time the effective number of nodes ($N_e$) and edges ($E_e$).

For the sake of simplicity, we will assume that the invasion period $1/\lambda$ is an integer. Let $P(t)$ be a row vector such that $(P(t))_l$ is the probability that the population contains $l$ distinct species at time $t$. We have

$$P(t) = P(0)\,\mathbf{G}^{\lfloor \lambda t \rfloor}$$

with $P(0) = [1\,0\,\cdots\,0]$ and with the transition matrix

$$\mathbf{G} = \begin{bmatrix} \frac{1}{S} & \frac{S-1}{S} & 0 & ... & 0 \\ 0 & \frac{2}{S} & \frac{S-2}{S} & ... & 0 \\ 0 & 0 & \frac{3}{S} & ... & 0 \\ ... & ... & ... & ... & ... \\ 0 & 0 & 0 & ... & 1 \end{bmatrix}.$$

In particular, we have

$$(P(t))_l = P(0)\,\mathbf{G}^{\lfloor \lambda t \rfloor} e_l$$

where $e_l$ is the $l$-th canonical vector of $\mathbb{R}^S$ (i.e. $e_l = [0\,\cdots\,1\,\cdots\,0]^T$ where the $l$-th component is equal to one).

**Nodes.** The average number of times the population contains exactly $l$ species over the time interval $[0, t]$ is given by

$$\sum_{\tau=0}^{\lfloor \lambda t \rfloor - 1} \frac{1}{\lambda} P(0)\,\mathbf{G}^{\tau} e_l + R(t) = \frac{1}{\lambda} P(0)(\mathbf{I} - \mathbf{G})^{-1}\left(\mathbf{I} - \mathbf{G}^{\lfloor \lambda t \rfloor}\right) e_l + R(t).$$

where $R(t) = \left(t - \frac{\lfloor \lambda t \rfloor}{\lambda} + 1\right) P(0)\,\mathbf{G}^{\lfloor \lambda t \rfloor} e_l$ is the remaining number of counts during the last incomplete period. Since $R(t)/t$ tends to $0$ as $t$ tends to infinity, this term can be discarded for $t$ large enough. It follows that the node associated with $l$ species has a weight equal to

$$\frac{1}{\lambda t} P(0)(\mathbf{I} - \mathbf{G})^{-1}\left(\mathbf{I} - \mathbf{G}^{\lfloor \lambda t \rfloor}\right) e_l + \frac{R(t)}{t}.$$

Note that this weight can be interpreted as the probability to observe exactly $l$ species at a uniformly randomly chosen time over the interval $[0, t]$. Finally, the weight of a node associated with a specific population containing $l$ species is given by

$$P_N = \frac{1}{\binom{S}{l}} \frac{1}{\lambda t} P(0)(\mathbf{I} - \mathbf{G})^{-1}\left(\mathbf{I} - \mathbf{G}^{\lfloor \lambda t \rfloor}\right) e_l + \frac{R(t)}{t}.$$

**Edges.** Since the graph is acyclic, the transition from level $l$ to level $l+1$ can be observed only once. Therefore, the average number of transitions from $l$ species to $l+1$ species over the interval $[0,t]$ is given by

$$\sum_{s=l+1}^{S}(P(t))_s = P(0)\,\mathbf{G}^{\lfloor\lambda t\rfloor}g_l$$

where $g_l = [0\,\cdots\,0\,1\,\cdots\,1]^T$ is a vector whose first $l$ components are zero and last $S-l$ components are equal to one. It follows that the probability of observing the transition from $l$ species to $l+1$ species at a uniformly randomly chosen time over the interval $[0,t]$ is equal to

$$\frac{1}{t}P(0)\,\mathbf{G}^{\lfloor\lambda t\rfloor}g_l.$$

This could be associated with the weight of the corresponding edge of the graph. Instead, the probability to observe the transition from $l$ species to $l+1$ species *among all randomly chosen transitions* that occurred over the time interval $[0,t]$ is given by

$$\sum_{s=l+1}^{S}\frac{1}{s-1}(P(t))_s = P(0)\,\mathbf{G}^{\lfloor\lambda t\rfloor}h_l$$

where $h_l = \left[0\,\cdots\,0\,\frac{1}{l}\,\frac{1}{l+1}\,\cdots\,\frac{1}{S-1}\right]^T$ is a vector whose first $l$ components are zero. Moreover, the number of edges from all populations of $l$ species to all populations of $l+1$ species is equal to $\binom{S}{l}\cdot(S-l)$. Then the weight of an edge from a specific population of $l$ species to a specific population of $l+1$ species is

$$P_E = \frac{1}{\binom{S}{l}\cdot(S-l)}P(0)\,\mathbf{G}^{\lfloor\lambda t\rfloor}h_l.$$

However, these values are only valid for a number of replicates tending to $\infty$. For a limited number of replicates, due to the sampling effect, some nodes and edges within the same $l$ may be observed more or less often than expected, resulting in larger differences in their observed proportions, especially in high $S$. These differences translate into larger differences in weights and therefore lower $N_e$ and $E_e$. For this reason, we also simulated null TIGs with the same number of replicates as the other TIGs.

### B Maximal values comparison plot

By observing null TIGs and simulated TIGS (with interspecific interactions)$N_e$ and $E_e$ over time, we observed that assembly was first random in a expanding phase leading to in increase of both metrics before reaching a maximal point. After it, a contracting phase was ruled by the interactions, inducing a decrease of both metrics. We thus wondered if the maximal value for $N_e$ and $E_e$ were temporaly correlated.

For each of the 500 main simulated TIGs, we divided $N_e$ and $E_e$ values by their maxima. These new metrics are thus the proportional values of $N_e$ and $E_e$ towards their maximum. To determine whether $max(N_e)$ and $max(E_e)$ co-occur, we examined each proportional values at time step when the other metric reached its maximum. In other words, we observed $\frac{N_e}{max(N_e)}$ when $E_e = max(E_e)$ (left of the plot) and $\frac{E_e}{max(E_e)}$ when $N_e = max(N_e)$ (right of the plot).

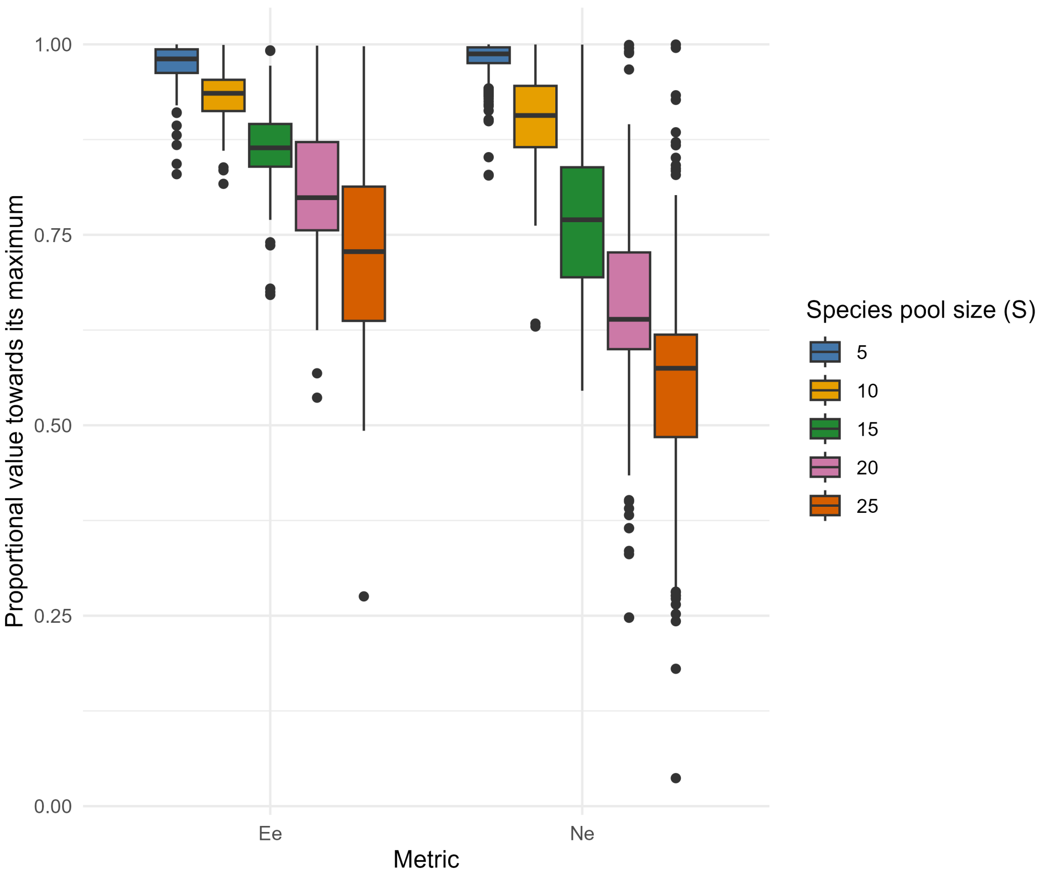


We observed that although $max(N_e)$ and $max(E_e)$ are usually close, they do not always co-occur, especially when $S$ is high. Indeed, a high $S$ induces a higher bound for $N_e$ and $E_e$, increasing the differences. That is why $max(N_e)$ and $max(E_e)$ are interpreted as boundaries of $N_e$ and $E_e$ diversities rather than as describing an actual TIG at a given time step.

### C Gini-like area index plot

In the contracting phase, both $N_e$ and $E_e$ decreased. However, not at the same rate, inducing a curvature in the $E_e \sim N_e$ trajectory. To study this curvature, we used the Gini-like area index returning a value between $-0.5$ (convex, $\bigcup$ shape and $0.5$ (concave, $\bigcap$ shape).

For each contracting phase, we normalized $N_e$ and $E_e$ to fit them in a range between $0$ and $1$. We next applied the Gini-like area index, which determines the curvature of the relationship. A negative coefficient indicates a convex curvature and a positive one indicates a concave curvature.

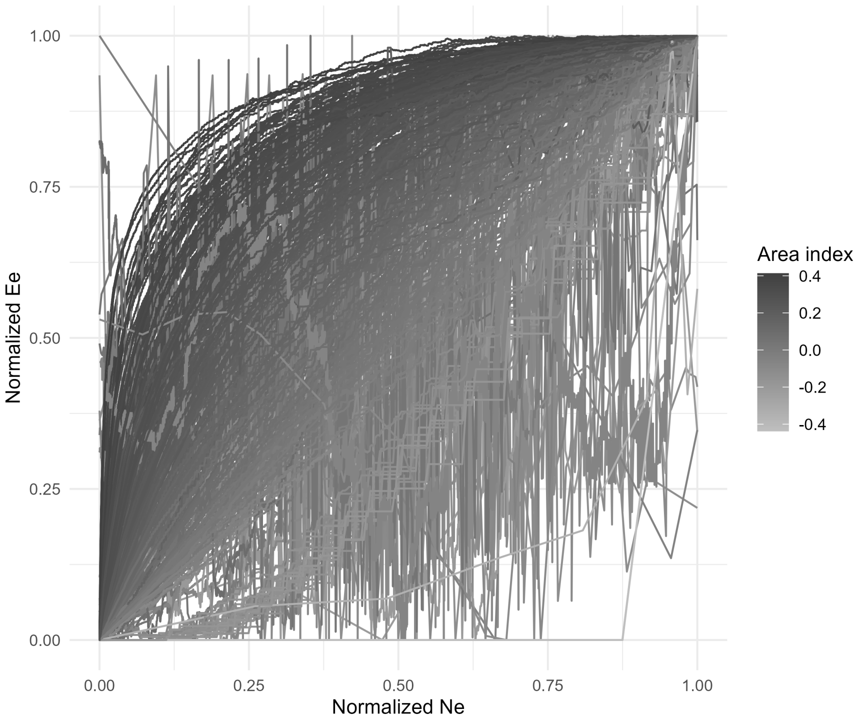


This plot represents every normalized contracting phase with a color according to the area index. For most TIGs, the area index is a strong indicator of the curvature gradient.

Some TIGs contain strong equilibria: once these are reached, no further transitions to other communities occur (transition probability at 0). In these cases, edges gain no more weights and $E_e$ stabilizes at a high value. The contracting phase is thus flattened and slightly convex. But with the normalization, the convex shape is stronger and the area index lower. That is why TIGs with a transition probability of 0 during the second half of the time were removed in the main text.

### D Repeated invasions detail

At the end of the observation times, $N_e$ was related to the number of uninvadable equilibrium. However, in a TIG context, repeated invasions can induce an invasion on such an equilibrium, and induce a transition to another equilibrium.

When the system is at an uninvadable equilibrium, every invasion growth rate ($IGRs$) (growth rate of an invader at the beginning of the invasion) are negative (no species can invade). However, some species can have an $IGR$ close enough to 0 that they do not reach $N = 0$ during the 10 steps between two invasions. In this case, if this species is selected twice in a row for an invasion attempt, its density will go above the detection threshold and the community will be temporarily the original community plus an invader. Because successive invasion attempts by the same species are more likely when $\frac{\lambda}{S}$ is high, this ratio determines the frequency of this phenomenon.

In some TIGs, such as the one below the line in Figure 4, this pattern can lead to a transition inducing a low occurrence of a stable community. Indeed, if an invader has an $IGR$ very close or equal to 0, it can be selected many time before vanishing, leading to high densities and a transition to another equilibrium.

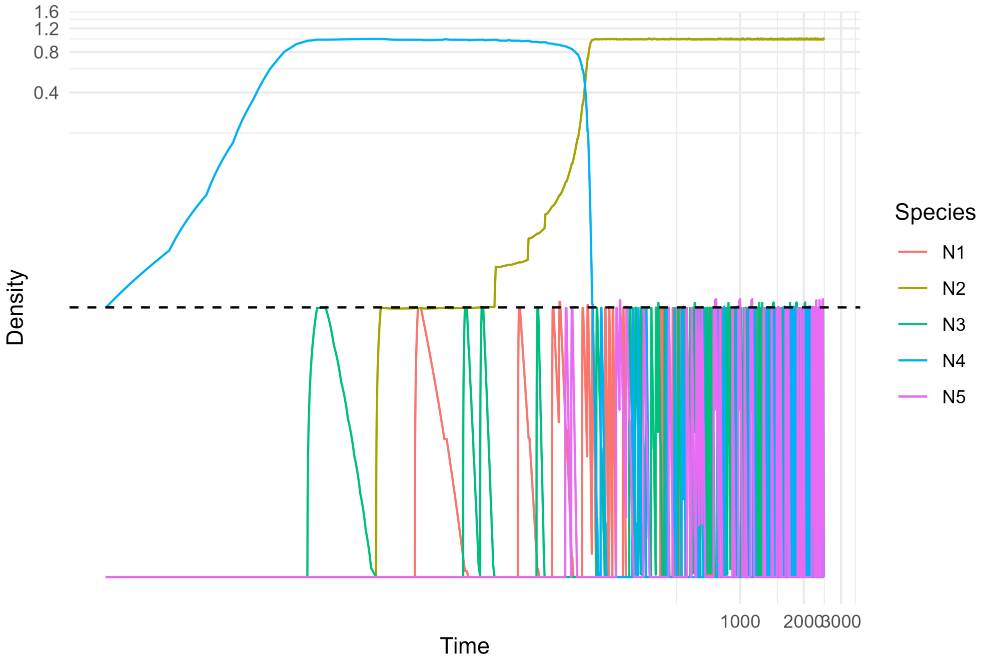


In this simulation, communities $\{2\}$ and $\{4\}$ each contain an uninvadable equilibrium. When the community is $\{4\}$, the second species' $IGR$ is negative but close to $0$. When this second species first invades, it stays below the detection threshold (dashed line) but does not go extinct quickly, so, during a second invasion attempt, the density goes above the detection threshold and is high enough to allow the second species to grow (the community is thus $\{2{,}4\}$. At a point (and after more invasions from the second species), the fourth species goes under the detection threshold, leading to the stable community $\{2\}$. $\{2\}$ and $\{4\}$ are both stable communities, but due to this pattern and repeated invasions, $\{4\}$ will always be replaced, affecting $N_e$ for $t \to \infty$.

### E Levels in canonical communities

We last applied TIG concept to four canonical coomunities with distinct dynamics: (1) a single and globally stable equilibrium, (2) multiple equilibria, depending on invasion history, (3) cycling, and (4) chaos. With them, we observed their median level and the variations around it all along the observation time.

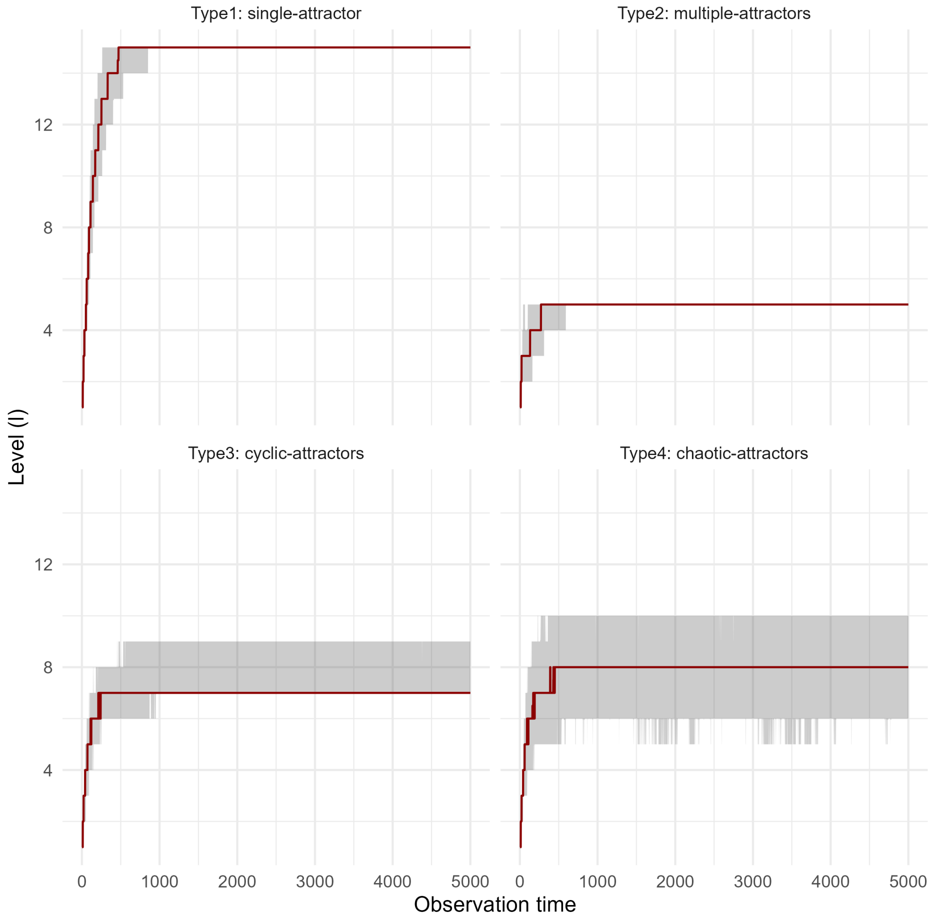


This figure shows the median level (red line) and its $90\%$ interval (grey area) of the four canonical communities over time. It shows how type 1 reached $l = 15$, while type 2 never went over $l = 7$, with types 3 and 4 at similar levels.

### F Simulation metrics

All of our results came from *in silico* simulations. They were made from repeated invasions from randomly selected species during an observation time ($t$) across several replicates. However, the required $t$ and number of replicates were determined by the species pool size $S$. All details are contained in the following table (with **Rep** as the number of replicates).

| **S** | **t** | **Rep** |
|---|---|---|
| 5 | 2500 | 300 |
| 10 | 3750 | 350 |
| 15 | 5000 | 400 |
| 20 | 6250 | 450 |
| 25 | 7500 | 500 |
